# Phospho-regulation of Atoh1 is required for plasticity of secretory progenitors and tissue regeneration

**DOI:** 10.1101/268607

**Authors:** Goran Tomic, Edward Morrissey, Sarah Kozar, Shani Ben-Moshe, Alice Hoyle, Roberta Azzarelli, Richard Kemp, Chandra Sekhar Reddy Chilamakuri, Shalev Itzkovitz, Anna Philpott, Douglas J. Winton

## Abstract

The intestinal epithelium is maintained by a small number of self-renewing stem cells in homeostasis. In addition committed progenitors can contribute to the functional stem cell compartment at a low level during homeostasis and substantially during regeneration following tissue damage. However the mechanism of, and requirement for, progenitor plasticity in mediating pathological response has not been demonstrated. Here we show that multisite phosphorylation of the transcription factor Atoh1 is required both for the contribution of secretory progenitors to the intestinal stem cell pool and for a robust regenerative response following damage. In lineage tracing experiments Atoh1^+^ cells (*Atoh1*^*(WT)CreERT2*^ mice) show stem cell activity by giving rise to multilineage intestinal clones both in the steady state and after tissue damage. Notably in the colonic epithelium a single generation of Atoh1^+^ progenitors sustains 1 in 15 stem cells. In an activating *Atoh1*^*(9S/T-A)CreERT2*^ line, the loss of phosphorylation sites on the Atoh1 protein promotes secretory differentiation and inhibits the contribution of these cells to self-renewal. Finally, in a chemical colitis model the Atoh1^+^ cells of *Atoh1*^*(9S/T-A)CreERT2*^ mice have reduced clonogenic capacity that impacts overall regenerative response of the epithelium. Thus progenitor plasticity plays an integral part in maintaining robust self-renewal in the intestinal epithelium and the balance between stem and progenitor fate behaviour is directly co-ordinated by Atoh1 multi-site phosphorylation.

## INTRODUCTION

Within the intestinal epithelium, cell generation occurs from phenotypically heterogenous stem cells residing at the base of glandular crypts (Vermeulen and Snippert, 2014). There is broad consensus that this heterogeneity reflects the combined behaviour of active and reserve stem cells. The former dominates in homeostatic self-renewal and the latter following tissue damage. In homeostasis, rapidly cycling stem cells express the R-spondin receptor Lgr5. Reserve stem cell function is less defined and has been ascribed variously to: a subset of quiescent Lgr5^+^ cells (Barriga et al., 2017; Buczacki et al., 2013); progenitors committed to different intestinal lineages (van Es et al., 2012; Tetteh et al., 2016); cells dependent on alternate pathways for stem cell maintenance (Takeda et al., 2011; Tian et al., 2011).

Previously it has been demonstrated that secretory progenitors possess reserve stem cell function in the small intestinal (SI) epithelium following tissue damage (van Es et al., 2012; Yan et al., 2017; Ishibashi et al., 2018). Subsequent to Delta-like expression (from *Dll1* or *Dll4*) the bHLH transcription factor Atoh1 is upregulated, an event required for the creation of all secretory lineages within the epithelium (Yang et al., 2001). Atoh1 can be phosphorylated on multiple sites by cyclin-dependent kinases. Here we demonstrate that maintenance of the plasticity of committed secretory precursors allowing return to the stem compartment is dependent on the multi-site phosphorylation of Atoh1, prevention of which inhibits Atoh1-mediated self-renewal and results in compromised regeneration following damage. We conclude that reversibility in commitment to differentiate is dependent on post-translational control of Atoh1 and is required to maintain a robust stem cell population.

## RESULTS

### Atoh1^+^ cells show stem cell activity

Initially to determine the extent to which Atoh1 positive cells support stem cell maintenance in homeostasis, we generated a mouse (*Atoh1*^*(WT)CreERT2*^) with an inducible *CreER*^*T2*^ downstream of the *Atoh1* coding sequence (Figure S1A). Acute lineage tracing demonstrated that tdTomato (tdTom) reporter expression 24 h following a single pulse of tamoxifen was restricted to secretory cells within the SI and colonic epithelium (Figures 1A-D, Figures S1B-S1G). Mature Paneth and goblet cells were positive for reporter, while enteroendocrine cells (EEC) were not. However, by 4 days post-tamoxifen EEC cells were also labelled (Figure 1E). Individual Paneth cells remain labelled for many weeks reflecting their longevity (Figure S1H). Similar results were found in the colon and long-lived secretory cells were also identified (Figure S1I). By 30 days post-induction cohesive patches of reporter positive cells that occupied all or a significant portion of entire crypts were present (Figures 1F and 1G), and continued to be observed after several months (Figure S1J). Immunostaining established the presence of goblet, Paneth, enteroendocrine, and absorptive cells within reporter positive epithelium confirming their multilineage composition (Figures 1H-1K). These patterns are identical to those arising from individual marked intestinal stem cells (Vermeulen et al., 2013) and demonstrate a clonal origin from Atoh1^+^ precursors. *Atoh1*^*(WT)CreERT2*^*; Rosa26*^*TdTom*^ mice were then crossed onto *Lgr5*^*Gfp*^ reporter mice to investigate co-expression of Atoh1 and the intestinal stem cell marker Lgr5. The expression of Atoh1 and tdTom reporter was identified in between 2-4% of Lgr5^+^ (GFP^+^) cells (Figures S1K-S1O), representing a likely intermediate state in the commitment process and candidate clonogenic population. Together these results confirm that *Atoh1* is appropriately expressed in mature Paneth/goblet cell but not enteroendocrine cells and that a proportion of Atoh1^+^ progenitors are acting as long-term multipotential stem cells (Bjerknes et al., 2012; Sommer and Mostoslavsky, 2014; Ishibashi et al., 2018).

**Figure 1.**
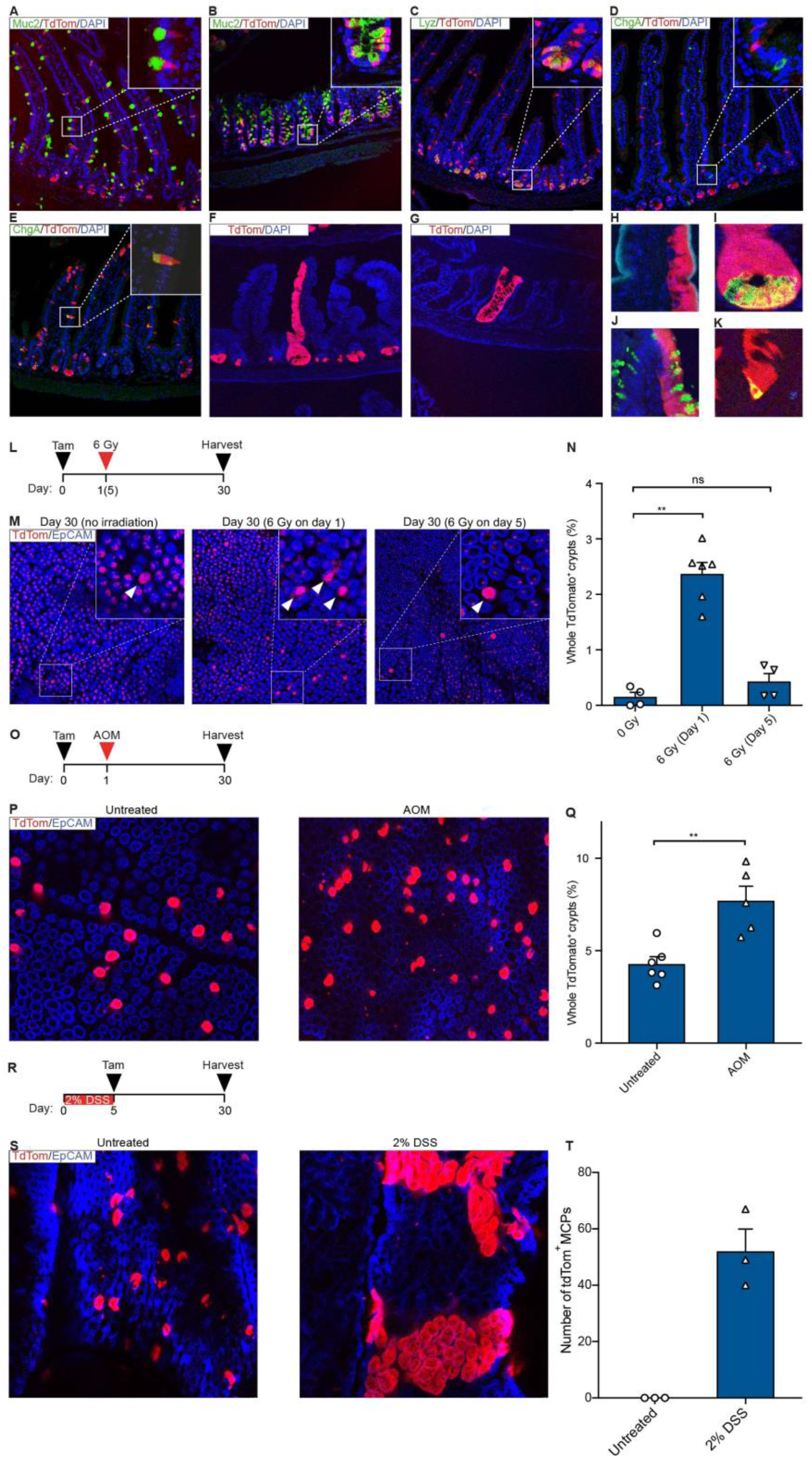
Lineage tracing of Atoh1^+^ cells in homeostasis and after injury. (A-D) tdTom reporter is detected in Muc2^+^ goblet cells in the SI (A), colon (B), and Lyz^+^ Paneth cells (C), but not in ChgA^+^ enteroendocrine cells 24 h post-tamoxifen (D). Muc2,Mucin2; Lyz, Lysozyme; ChgA, Chromogranin A. (E) ChgA^+^ cells labelled with tdTom at day 4 after induction. (F, G) Reporter positive clone in the SI (F), and colon (G), 30 days following tamoxifen. (H-K) tdTom^+^ clones at 30 days are composed of Alpi^+^ enterocytes (H), Paneth cells (I), goblet cells (J), and enteroendocrine cells (K). (L, M, N) Schematic of induction and injury protocol (irradiation (L), AOM (O), DSS (R)). (N) Representative pictures of SI whole-mounts containing labelled crypts (arrowheads) 30 days postinduction. (M) Quantification of tdTom^+^ crypts in the SI (n=4 for 0 Gy, n=6 for 6 Gy (Day 1), n=4 for 6 Gy (Day 5). (P, S) Representative images of colonic crypts at day 30 post-tamoxifen and AOM (P) or DSS treatment (S). Note large tdTom^+^ regenerative multi-crypt patches (MCPs) associated with 2% DSS treatment (S). (Q) Quantification of reporter positive crypts in colon (n=6 for untreated, n=5 for AOM treated). (T) Quantification of tdTom^+^ MCPs in untreated and DSS-treated colons (n=3 for both groups). The non-parametric Mann Whitney test was used (mean±s.e.m, ** P=0.0095 (C), ** P=0.0087 (F)). Alpi, Alkaline phosphatase. AOM, Azoxymethane; DSS, Dextran Sodium Sulphate. See also Figure S1.

### Atoh1^+^ cells contribute directly to epithelial regeneration

The extent of reversibility of Atoh1^+^ cell commitment was studied in the context of irradiation-induced tissue damage. Irradiation given 1 day after tamoxifen generated an increased number of tdTom^+^ crypts at 30 days in SI compared to unirradiated controls (16-fold increase, 2.37% vs. 0.15%). This effect was abrogated when irradiation was given 5 days after tamoxifen (Figures 1L-1N), suggesting that regenerative potential is a property of progenitors arising *de novo* from the stem cell compartment and not of more mature secretory cells. Similarly, after targeted deletion of the bulk of Lgr5^+^ stem cells using a diphtheria toxin approach there was a 30-fold increase in the number of clones observed (Figures S1P and S1Q). Adapting the assay to perform a similar analysis for the colonic epithelium and to circumvent that tissue’s known radio-resistance (Cai et al., 1997) mice were treated with the colon specific carcinogen azoxymethane (AOM) 1 day after tamoxifen treatment. Again, an increase in the frequency of tdTom^+^ crypts was observed (Figures 1O-1Q). Following dextran sodium sulphate (DSS)-induced colitis, multicrypt tdTom+ patches were detected at the margins of regions of damage (Figures 1R-1T; Figures S1R and S1S). Together these results suggest that Atoh1^+^ cells directly contribute to regeneration following damage.

### Creating a pro-secretory phosphomutant Atoh1

Previous studies have indicated that multi-site phosphorylation of bHLH proteins restrains cell cycle exit and limits differentiation, while conversely un(der)phosphorylation promotes these processes in the developing nervous system and pancreas (Ali et al., 2011, 2014; Azzarelli et al., 2017), although a role for multi-site phosphoregulation of bHLH proteins in adult homeostasis or tissue repair has not been reported. Hence, we hypothesised a potential role for Atoh1 phosphorylation in controlling transition between stem and progenitor compartments both in homeostasis and under conditions of heightened proliferation following tissue damage. Cyclin-dependent kinases phosphorylate on serine-proline (SP) or threonine-proline (TP) residues. Atoh1 has 9 S/T-P sites available for phosphorylation (Figures 2A-2C). The phosphorylation of two SP sites has previously been shown to destabilise the Atoh1 protein in the context of neuronal precursors (Forget et al., 2014). Atoh1 can be phosphorylated on many sites; we saw at least 5 distinct phospho-forms of Atoh1 after phosphorylation by different Cyclin/Cdk combinations (Figure 2B). We expressed forms of Atoh1 where S/T-P sites were mutated to alanine-proline (AP), in colorectal cancer cells to determine the effect of Atoh1 phosphorylation on cell proliferation and on expression of markers of differentiation. While mutation of the two phospho-sites previously described in neuronal precursors (Forget et al., 2014) had a modest effect on Atoh1 activity, mutation of all 9 S/T-P sites was more effective at promoting enhanced cell cycle exit (Figures 2D and 2E). Additionally, expression of secretory genes (Figure 2F) is enhanced after mutation of all 9 potential phosphorylation sites, compared to both wild-type Atoh1 and 2S/T-A Atoh1, consistent with multi-site phospho-regulation of Atoh1 playing a significant role in the controlling the balance between proliferation and differentiation (Ali et al., 2011; Azzarelli et al., 2017).

**Figure 2.**
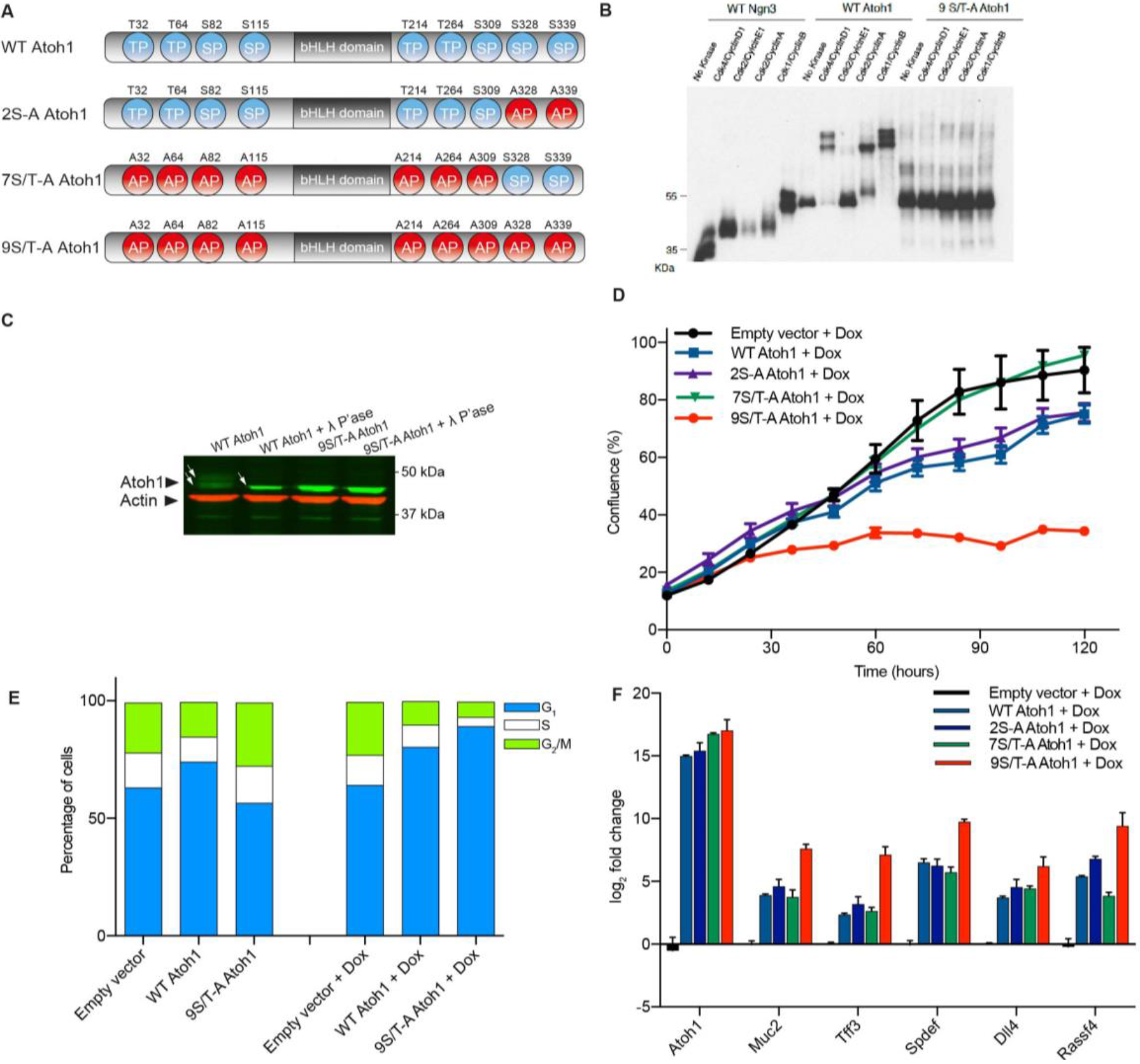
Identification of a hyperactive phosphomutant Atoh1. (A) Diagram depicting the location of proline-directed kinase motifs (SP or TP) in Atoh1 protein, and mutations of these sites into alanine in Atoh1 phosphomutants. (B) In vitro translated Atoh1 protein band-shift following incubation with different CDKs. Ngn3 was used as a positive control. (C) WT Atohl bands (arrows) collapse following λ phosphatase treatment, demonstrating phosphorylation. (D) Cell proliferation following Dox-induced expression of *WT* or phosphomutant *Atoh1* (n=3 biological replicates, 2 technical replicates). (E) Cell cycle profile of uninduced and Dox-treated cells showing increased G1 and decreased S/M populations on induction of *9S/T-A Atoh1*. (F) Gene expression of *Atoh1* and its target/secretory differentiation genes 72 h after Dox induction of DLD-1 cells (n=3 biological replicates, 2 technical replicates; *Gapdh*- normalised). SP, Serine-Proline; TP, Threonine-Proline; Dox, doxycycline. CDK, Cyclin-dependent kinase

### 9S/T-A phosphomutant Atoh1 promotes secretory maturation in vivo

To investigate how preventing phosphorylation of Atoh1 affects progenitor-mediated self-renewal in homeostasis and repair, we substituted 9S/T-A Atoh1 for the wild-type form in its endogenous locus, generating a knock-in mouse identical in design to
*Atoh1*^*(WT)CreERT2*^ but with the hyperactive phosphomutant *Atoh1*^*(9S/T-A)CreERT2*^ allele (Figure S2A). Heterozygous *Atoh1*^*(9S/T-A)CreERT2*/+^ and control *Atoh1*^*(WT)CreERT2*/+^ mice were crossed to establish homozygous *Atoh1*^*(WT)CreERT2*^ and *Atoh1*^*(9S/T-A)CreERT2*^ lines respectively. Phenotype analysis identified no gross differences between the two lines. Mice developed normally and the overall morphological appearance of the epithelium remained unchanged. More detailed analysis found no difference in the number or distribution of the different secretory lineages or in the frequency of apoptotic cells (Figures S2B-S2F).

To investigate if the 9S/T-A mutations impacted on secretory maturation after lineage specification, transcriptional profiling of secretory cells in the two lines was performed. First the expression profile of Atoh1^+^ cells from *Atoh1*^*(WT)CreERT2*^ mice was determined by comparing tdTom^+^ (secretory) and tdTom^-^ (absorptive) cells to define the baseline pro-secretory signature for both colon and small intestine (Table S1). Next the transcription profiles of tdTom^+^ cells from wildtype and mutant mice (Table S2) were compared against this Atoh1^+^ baseline and a published secretory signature (Lo et al., 2017). These gene set enrichment analyses (GSEA) demonstrated a major pro-secretory shift in *Atoh1*^*(9S/T-A)CreERT2*^ mice in both tissues and a strongly reduced intestinal stem cell signature compared to those from controls (Figures 3A-3E).

**Figure 3.**
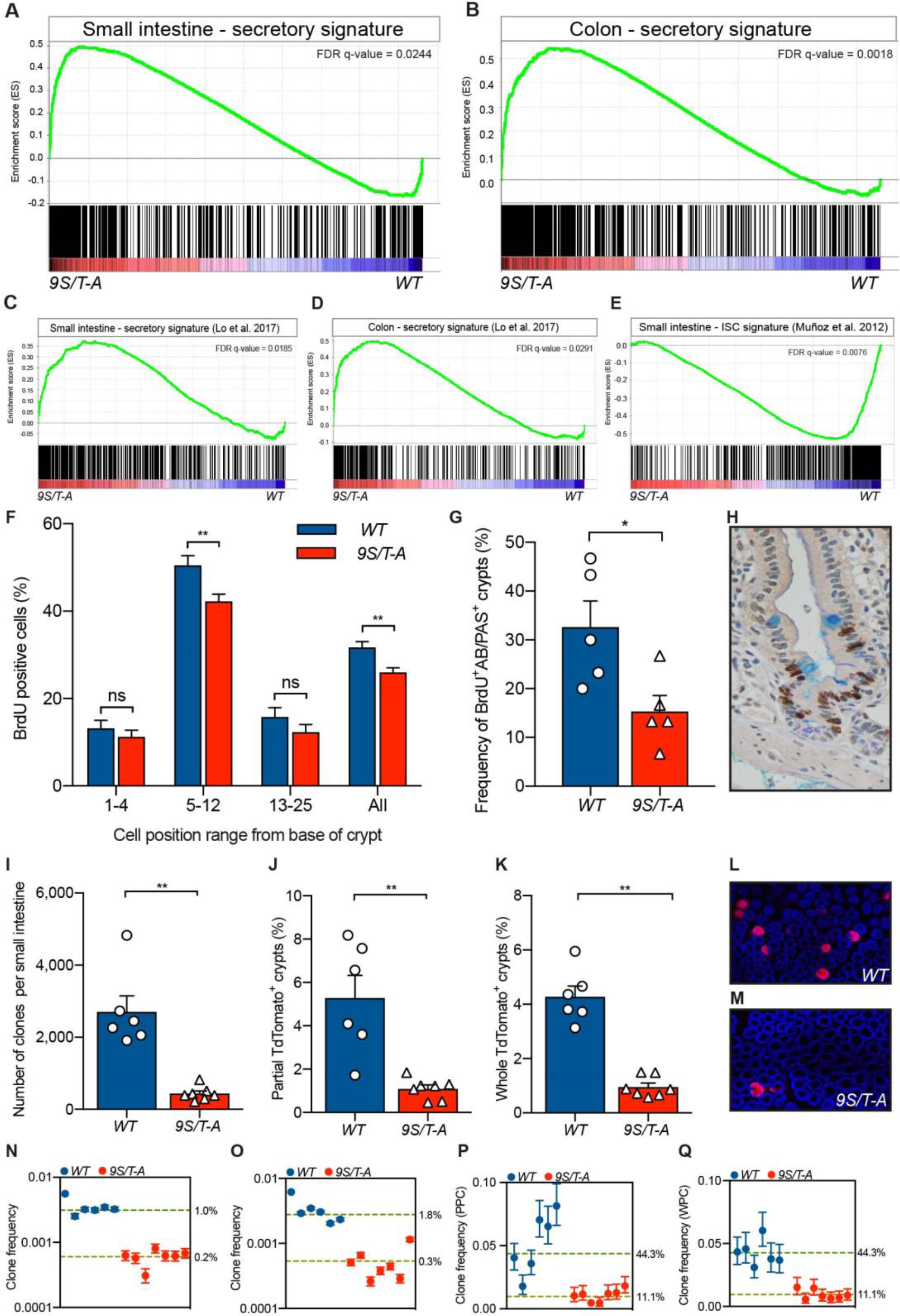
9S/T-A Atoh1 promotes secretory maturation and reduces proliferation and the number of clonogenic Atoh1^+^ cells. (A, B) Gene Set Enrichment Analysis (GSEA) of Atoh1^+^ SI secretory signature (A) and colon (B) shows an enrichment of secretory genes in 9S/T-A Atoh1 tdTom^+^ cells. (C, D) GSEA utilising a published secretory transcriptome reveals an increase in secretory gene signature in phosphomutant-expressing tdTom^+^ cells in SI (C) and colon (D). (E) GSEA using a published intestinal stem cell (ISC) gene signature (Muñoz et al., 2012) shows a de-enrichment of ISC genes in the mutant SI progenitors. (F-G) BrdU labelling index for a range of cell positions in SI crypt (F) shows a reduction in proliferation above the crypt base (n=100 crypts, 4 mice per genotype; mean±s.e.m; ** P=0.0061 and 0.0015). (F) Frequency of crypt-villus units containing at least one BrdU^+^ goblet cell after a 24 h BrdU-pulse (n=5 for both groups, * P=0.0317). (H) Representative image of a crypt-villus unit with a BrdU^+^AB/PAS^+^ cell. (I) Quantification of tdTom^+^ clonal events in the SI (n=6 (*WT*), n=7 (*9S/T-A*), mean±s.e.m, ** P=0.0012). (J) Partly populated tdTom^+^ crypts in colon (n=6 (WT), n=7 (*9S/T-A*), mean±s.e.m, ** P=0.0023). (K) Whole tdTom^+^ crypts in *WT* and *9S/T-A* colons (n=6 (*WT*), n=7 (*9S/T-A*), mean±s.e.m, ** P=0.0012; Same *WT* data was shown in Figure 1Q as the experiment was done in parallel). All samples were collected 30 days after tamoxifen. Mann Whitney test was used for all comparisons. (L, M) Representative images of *WT* (L) and *9S/T-A* (M) colons scored in J and K. (N-Q) Inference of the proportion of the clonogenic fraction of labelled Atoh1^+^ cells in proximal SI (N), distal SI (O), colon-PPC (P), and colon- WPC (Q). The numbers next to the dotted lines indicate the inferred proportion of crypts that had one labelled stem cell. AB/PAS, Alcian Blue/Periodic Acid Schiff.

The pro-secretory nature of 9S/T-A expressing cells arose from an overall elevation in pro-secretory transcripts for goblet and Paneth cell lineages (SI only). GSEA identified that SI Atoh1^+^ cells isolated from *Atoh1*^*(9S/T-A)CreERT2*^ mice had a reduction in a subset of transcripts associated with EECs, indicating that Atoh1 phosphorylation influences their maturation (Figures S3A and S3B).

### Epithelial proliferation and clonogenicity are inhibited in *Atoh1*^*9S/T-A*^ mice

Our GSEA analyses indicate that preventing multi-site phosphorylation of Atoh1 enhances pro-secretory gene expression. We next investigated whether this was accompanied by changes in proliferation. Comparing proliferation between the two lines demonstrated a slight overall decrease in the total proliferative index of the crypts of *Atoh1*^*(9S/T-A)CreERT2*^ mice in both SI and colon but that did not reach significance in the latter. More detailed spatial analysis within the crypt epithelium demonstrated that this effect was largely accounted for by a decrease in the proportion of cells in S-phase in the epithelium of *9S/T-A* mutants in cell positions above the very base of the crypt and a reduction in the frequency of proliferative goblet cells (Figures 3F-3H; Figure S2G). This supports the interpretation that the phosphorylation of Atoh1 in cells immediately arising from the stem cell population limits Atoh1-dependent cell cycle exit to allow maintenance of proliferation in progenitors, while preventing this phosphorylation limits the ability to return to a proliferative stem/progenitor compartment. We next tested this hypothesis using a lineage tracing approach.

Lineage tracing and FACS analysis established that the acute pattern of reporter expression and absolute number of tdTom^+^ cells was the same in *Atoh1*^*(9S/T-A)CreERT2*^ and controls (Figures S2H-S2O). However, lineage tracing at 30 days identified fewer epithelial clones in both SI and colon than in *Atoh1*^*(WT)CreERT2*^ mice (Figures 3I-3M). This demonstrates that preventing phosphorylation of Atoh1 impairs the return of Atoh1^+^ cells to the stem cell compartment, and confirms a role for Atoh1 phosphorylation in maintenance of progenitor plasticity.

Previously we and others have described that only a subset of competing stem cells drive increases in clone sizes that lead to surviving clones populating entire crypts (Kozar et al., 2013; Ritsma et al., 2014). To determine the net contribution of Atoh1^+^ cells to this population, mathematical modelling was used to infer the proportion of the clonogenic fraction that are initially marked in *Atoh1*^*(WT)CreERT2*^ and *Atoh1*^*(9S/T-A)CreERT2*^ mice (Supplementary information). In both the SI and colon the contribution of Atoh1 ^+^ progenitors to the stem cell pool is reduced in *9S/T-A* animals (Figures 3N-3Q). Between 1% and 2% of SI crypts in *Atoh1*^*(WT)CreERT2*^ mice contain a single clonogenic stem cell derived from an Atoh1^+^ progenitor and this is reduced five-fold in *Atoh1*^*(9S/T-A)CreERT2*^ mice (Figures 3N and 3O). In colon values are higher, with the observed 4% wholly populated crypts (WPC) and 5% partly populated crypts (PPC) identified in *Atoh1*^*(WT)CreERT2*^ mice at 30 days post-induction requiring that initially 44% of crypts (1 in 15 active stem cells) contained an Atoh1^+^ derived stem cell. This is reduced to 11% in *9S/T-A* mutant mice (Figures 3P and 3Q). Notably these rates reflect the contribution of a single cohort of transient progenitors arising from the stem cell pool that are produced over one or two days.

### Compromised epithelial regeneration in *Atoh1*^*9S/T-A*^ mice

While phosphorylation of Atoh1 clearly regulates reversion of secretory progenitors to the stem cell compartment, the absence of any other apparent phenotype in the 9S/T-A Atoh1 mice suggests a limited requirement for such plasticity in homeostasis. We next investigated the role of Atoh1 phosphorylation in mounting a robust regenerative response following tissue damage. In the DSS-induced chemical colitis model, *9S/T-A* mutant mice showed a greater sensitivity after treatment with increased weight loss and slowed recovery (Figures 4A, S4A, and S4B). Analysis of this phenotype at the start of the regenerative phase (9 days after the start of DSS treatment) showed areas of ulceration that were more extensive in mice carrying the *9S/T-A* mutant (Figures 4B and 4C). At both 5 and 9 days the proportion of secretory versus epithelial cells is identical for the two lines and cell death was restricted to a few cells on the luminal surface suggesting that the greater sensitivity does not arise from enhanced damage or deletion of secretory cells in *9S/T-A* mutant mice (Figures 4D-4F). However lineage tracing at 30 days following DSS treatment identified a reduced number and size of tdTom^+^ regenerative patches in *9S/T-A* colons compared with *WT* (Figures 4G, 4H, S4C, and S4D). Together these results demonstrate that mice lacking the ability to phospho-regulate Atoh1 have compromised regenerative capacity following damage and that the contribution of Atoh1^+^ progenitors is required for robust tissue repair.

**Figure 4.**
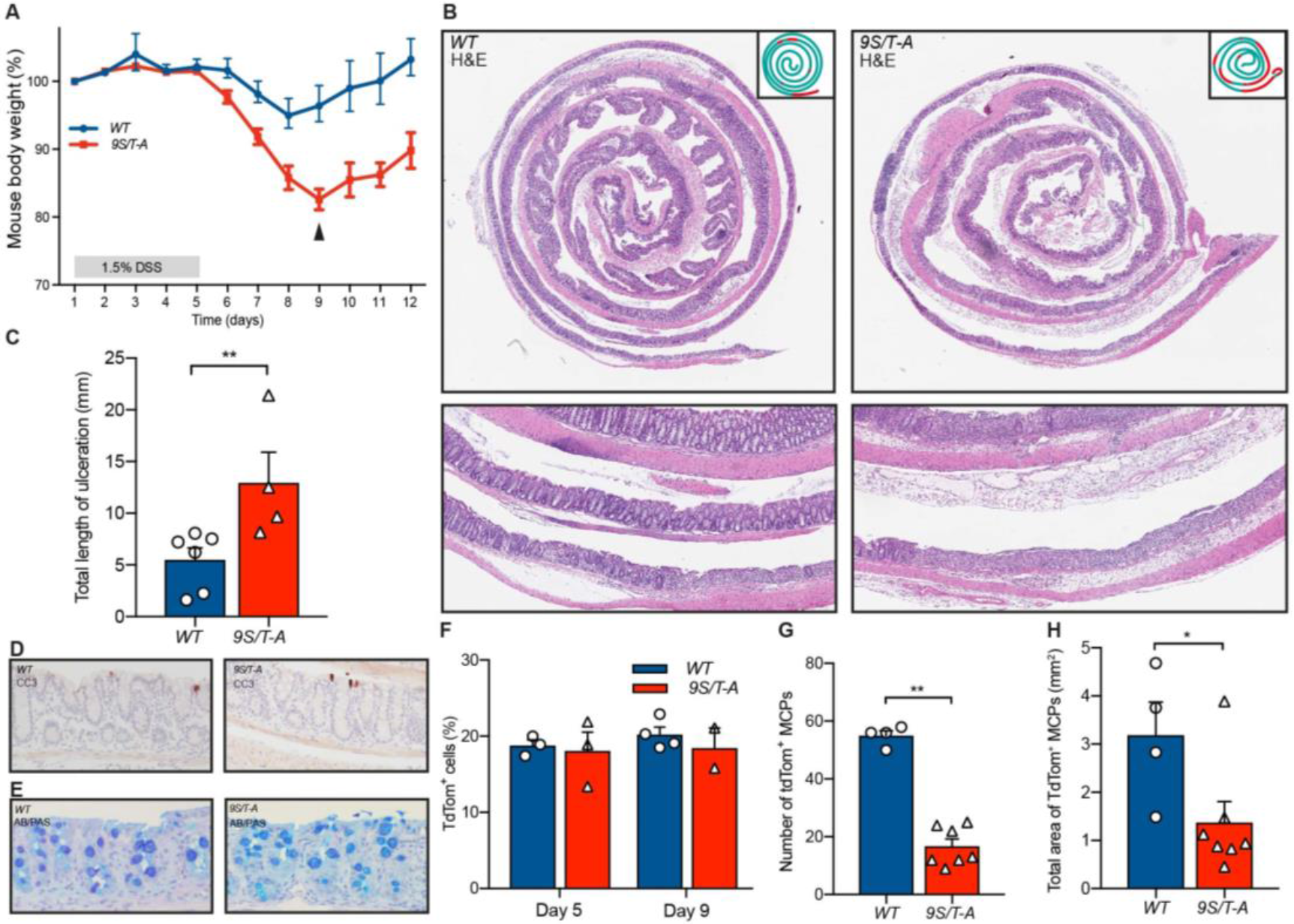
*Atoh1*^*(9S/T-A)CreERT2*^ mice are sensitive to chemical colitis. (A) Change in mouse body weight during and after DSS treatment (n=5 (*WT*), n=6 (*9S/T-A*); two *9S/T-A* mice were euthanized on day 9 on health grounds, and one *WT* animal was taken for a comparison (arrowhead)). (B) Representative pictures and schematics of colon on day 9 showing extensive loss of crypts in *9S/T-A*, but not in *WT*. (C) Total length of colon ulceration on day 9 (n=6 (*WT*), n=4 (*9S/T-A*), mean±s.e.m, ** P=0.0095). (D, E) Representative images of distal colon on day 5 of DSS treatment showing apoptosis (D) and AB/PAS staining (E) in *WT* and *9S/T-A* animals. (F) FACS analysis of the number of tdTom^+^ cells during DSS-induced colitis. (G, H) Analysis of the number (G) and the total area (H) of tdTom^+^ multi-crypt patches (MCPs) following 1.5% DSS (n=4 (*WT*), n=7 (*9S/T-A*), mean±s.e.m, ** P=0.0061, * P= 0.0424). AB/PAS, Alcian Blue/Periodic Acid Schiff. See also Figure S4.

## DISCUSSION

It is now accepted that cells with the capacity for self-renewal arise from a larger population whose members all have the same potential for self-renewal subject to occupying available niches (Farin et al., 2016; Ritsma et al., 2014). Here we show that Atoh1^+^ cells make a more substantial contribution to stem cell maintenance from cells committing to secretory differentiation than has been recognised hitherto (van Es et al., 2012; Ishibashi et al., 2018). Self-renewal is therefore not solely a feature driven from a fixed pool of stem cells but rather involves dynamic interchange between progenitors and stem cells in the steady state.

Transcription factors of the bHLH family have been extensively studied as master regulators of cell fate commitment and differentiation in a very wide variety of tissues including the nervous system and intestine (Ali et al., 2011, 2014; Yang et al., 2001). However, in recent years additional roles for these proteins are emerging in direct co-ordination of cell cycle and differentiation events particularly during embryonic development (Castro and Guillemot, 2011). Intestinal homeostasis in many ways represents an ongoing development-like hierarchical process, where crypts are maintained by stem cells feeding a proliferating progenitor compartment that gives rise to a variety of mature cell types. What is now also emerging is a picture of significant plasticity, where cells expressing Atoh1, previously thought to represent a population that have undergone secretory commitment, can nevertheless revert to “stemness” and repopulate the entire crypt with surprisingly high frequency. Mechanisms controlling this plasticity have been unclear. We see that the balance between stem and progenitor fate behaviour in the intestine can be controlled by Atoh1 multi-site phosphorylation under normal homeostatic conditions.

Control of proliferation and differentiation by modulation of bHLH protein phosphorylation is emerging as an important mechanism in development of the nervous system and the pancreas (Cleaver, 2017; Guillemot and Hassan, 2017). We now see that multi-site phosphorylation is also required to restrain irreversible commitment of secretory precursors in the adult homeostatic gut and so to maintain their ability to repopulate the stem cell compartment. Consistent with this, a phosphomutant form of Atoh1 enhances expression of gene sets associated with a more mature secretory phenotype in colorectal carcinoma cells. Interestingly, in the normal homeostatic gut despite Atoh1^+^ cells normally supplying up to 1 in 15 cells in the stem cell compartment, the phosphomutant Atoh1-expressing intestine is essentially phenotypically normal, indicating that plasticity from the secretory to stem compartment is not essential in normal homeostasis. However, intestinal regeneration after damage is substantially compromised by an inability to phosphorylate Atoh1.

Taken together, our results indicate that multi-site phosphorylation of Atoh1 is used to dynamically regulate the ability of secretory precursors to return to the stem cell compartment, in particular facilitating the capacity to respond rapidly to changes in the cellular environment. Damaging the intestine using irradiation or DSS (van Es et al., 2012; Ishibashi et al., 2018) leads to acute cell damage and death, followed by proliferative regeneration that produces new cells for tissue repair. Activation of CDKs and MAP kinases in rapidly proliferating cells undergoing regeneration would result in enhanced phosphorylation of Atoh1 restraining further progression down the secretory lineage and supporting re-entry of Atoh1-expressing cells into a stem-like state to allow rapid tissue regeneration. Thus, post-translational regulation of Atoh1 by proline-directed kinases to modulate the balance between proliferation and differentiation in response to changing tissue demands in the adult echoes the use of post-translational modification of other proneural proteins to changes the balance between proliferation and differentiation as development progresses (Hardwick et al., 2015). Progenitor plasticity is not merely an incidental acquired behaviour following damage but plays an integral part in tissue restoration and requires post-translational regulation of Atoh1.

## ACKNOWLEDGMENTS

This work was supported by Cancer Research UK (G.T., E.M., A.H., R.K., and D.J.W.), and Wellcome Trust Grant [103805] (G.T., D.J.W.). A.P. was funded by MRC Research Grant MR/K018329/1, Grant MR/L021129/1, Rosetrees Trust, Stoneygate Trust, and received core support from Wellcome Trust and MRC Cambridge Stem Cell Institute. We thank S. Taylor for DLD-1 Flp-In T-Rex cells, and D. Perera for plasmids. We also thank CRUK CI Core facilities and members of Winton and Philpott labs for their help.

## AUTHOR CONTRIBUTIONS

D.J.W., A.P., R.K., and G.T. designed the study. G.T. performed experiments. E.M. performed quantitative modelling. S.K., S.B-M., A.H., and R.A. performed some of the experiments. C.S.R.C. performed bioinformatics analyses. D.J.W. A.P., and G.T. wrote the manuscript with comments from all authors.

## METHODS

### Cloning of mouse knock-in constructs

For generation of mouse knock-ins *Atoh1* locus and homology arms were amplified from a bacterial artificial chromosome (BAC) RP24-77K22 (BACPAC Resources Centre). The targeting construct was assembled by a combination of seamless cloning (In-Fusion, Clontech) and restriction digest and ligation. For this a loxP site was introduced into 5’UTR of Atoh1 via PCR amplification. A neomycin cassette was inserted such that the 3’UTR was not disrupted. The CreER^T2^-hCD2-3’UTR was generated via gene synthesis service (Integrated DNA Technologies). The *Atoh1* sequence (*Atohl*^*(WT)*^ or *Atoh1*^(*9S/T-A*)^ was merged with this construct, and then ligated with Atoh1 vector containing the homology arms. The targeting vector sequence was verified by Sanger sequencing and linearised by SwaI enzyme before transfecting into ES cells. The final inserted sequence is available on request.

### ES cell targeting

Electroporation of the targeting construct into mouse ES cells was conducted by the CRUK CI Transgenic Core. ES cells were positively selected with G418. Correct integration of the construct was verified by long range PCR (SequalPrep, Thermo Fisher) according to the manufacturer’s instructions. Left integration arm was detected using a forward primer 5’-GGA CAG GCG GGA ACC ACA GA-3’ and a reverse primer 5’-TTG TCA ACA CGA GCT GGT CGA A-3’. Right integration arm was amplified using the following set of primers: forward 5’- CAA CAC AAC CCT GAC CTG TG-3’, and reverse 5’-CCC TAA CCA GTG TGC CCT TA-3’. Copy number of the clones was determined by qPCR of the neomycin selection cassette via a commercial genotyping service provider (Transnetyx). Single copy ES cell clones were taken forward for blastocyst injection, and chimeric mice were generated. Following successful germline transmission, the mice heterozygous for the targeting construct were crossed onto *PGK-Cre* line (Lallemand et al., 1998) in order to remove both the neo selection cassette and the endogenous *Atoh1* locus at the same time. A constitutively active *Atoh1-P2A-CreER*^*T2*^ allele was generated in this process.

### Mouse genotyping

Genotyping was carried out by Transnetyx. Manual genotyping by PCR was used to distinguish between homozygous and heterozygous Atoh1 animals. The following primers were used: forward 5’-TTT GTT GTT GTT GTT CGG GG-3’; reverse 5’-TCT TTT ACC TCA GCC CAC TCT T-3’.

### Creation of Doxycycline inducible DLD-1 cells

To generate an inducible stable cell line, a DLD-1 (human colon carcinoma) Flp-In T-Rex system-modified (Thermo Fisher) cell line containing a single Frt site was obtained (a generous gift from Prof Stephen Taylor, University of Manchester). The cell line authentication was carried out using Single Tandem Repeat (STR) genotyping. Tests were performed routinely to confirm mycoplasma-negative status of the cells. Atoh1 construct in a pcDNA 5/FRT/TO vector (Thermo Fisher) was co-transfected with pOG44 (Flp recombinase-expressing plasmid) in a 1:9 ratio (JetPrime, Polyplus transfection). Cells were washed 24 h after transfection, and fresh medium was added. Two days after transfection, the cells were split at a low confluence (less than 25%), and hygromycin (400 μg/mL) was added to the trypsinised cells. Fresh medium was added to the cells every 3-4 days, until the non-transfected cells died off, and foci of surviving cells could be visualised.

Validation of the correct recombination of the construct was carried out by PCR. Left integration arm was detected by using the following set of primers: forward 5’-AGT CAG CAA CCA TAG TCC CG-3’; reverse 5’- TTC TGC GGG CGA TTT GTG TA-3’. Correct integration on the 3’ end of the construct was done using a forward primer 5’-TAA ACG GCC ACA AGT TCA GC-3’, and a reverse primer 5’-CGG GCC TCT TCG CTA TTA CG-3’. Parental DLD-1 Flp-In T-Rex cell line was used as a negative control.

The expected loss of β-galactosidase activity on targeting was verified by X-Gal (5-bromo-4-chloro-3-indolyl-β-D-galactopyranoside) staining of fixed cells. To validate that the constructs were integrated as a single copy in the genome, copy number qPCR was employed. Copy number TaqMan probes detected HygR (Mr00661678_cn) and used a reference copy number assay for RNase P detection.

DLD-1 Flp-In T-Rex cells were maintained in Dulbecco’s Modified Eagle Medium (DMEM) with L-glutamine. Medium was supplemented with 10% Tet System-approved foetal bovine serum (FBS, Clontech). Doxycycline (100 ng/mL, Sigma) was added to the culture 24 h after seeding, to induce expression of the gene of interest. The cells were cultured under standard conditions.

### Treatment of animals

All animal experiments were carried out under the guidelines of the UK Home Office. Mice used in this study were 8-16 weeks old males and females of C57BL/6 background. Induction of CreER^T2^ in animals was carried out using the free base tamoxifen (Sigma) dissolved in ethanol/oil (1:9). The animals received 3 mg tamoxifen via an intra-peritoneal injection. To define Atoh1 secretory signature, the mice were injected with 1 mg tamoxifen per day on 3 consecutive days for a maximum labelling of all secretory lineages.

SI injury was induced by exposing animals to whole-body irradiation (6 Gy). To induce colon-specific injury, mice were given 1.5% DSS (MP Biomedicals) in drinking water for 5 days. DSS was replaced every two days during the treatment.

### Crypt fractionation and single cell preparation

SI (proximal 15 cm) and colon were dissected, flushed with PBS, everted and fed onto a glass rod spiral. They were incubated at 37°C in Hank’s Balanced Salt Solution (HBSS) without Ca^+2^ and Mg^+2^, containing 10 μM EDTA and 10 mM NaOH. Crypt release was facilitated using a vibrating stirrer (Chemap). Samples were incubated for 1 h and pulsed every 10 min. Fractions were collected after each pulse, and fresh solution added. Crypt-enriched fractions were pooled and washed in cold 2% FBS/PBS. Pooled fractions were resuspended in 0.05% trypsin and incubated for 7 min at 37°C, shaking every 1 min. Single cells were then filtered through a 70 μm mesh, and washed twice in 2% FBS/PBS.

### Flow cytometry

Single cell suspension obtained by trypsin treatment was washed and incubated with an anti-mouse CD326 (EpCAM) AlexaFluor 647 antibody (1:2,000, clone G8.8, Biolegend). DAPI (10 μg/mL) was added to distinguish between live and dead cells. Flow sorting was carried out on a BD FACS Aria SORP (BD Biosciences), using appropriate single-stained and unstained controls.

### Whole-mount preparation

Tissue was cut open, pinned out luminal side up, and fixed for 3 h at room temperature in ice-cold 4% PFA in PBS (pH 7.4). Whole-mounts were washed with PBS, and incubated with demucifying solution (3 mg/mL dithiothreitol (DTT), 20% ethanol, 10% glycerol, 0.6% NaCl, 10 mM Tris, pH 8.2) for 20 min, and mucus removed by washing with PBS.

### Whole-mount scanning and quantification

The tdTom fluorescence in colon whole-mounts was detected using Amersham Typhoon 5 laser scanner (GE Healthcare) at a 10 μm resolution. The tdTom^+^ foci were scored manually in Fiji. Mid and distal colon were scored only as the shape of the proximal colon prevented confident assessment of tdTom^+^ patches.

### Antibody staining

For staining whole-mount sections of 2 cm in length were excised, washed in 0.1% PBS-T for 2 days, and blocked in 10% donkey serum in PBS overnight at 4°C, protected from light. Samples were then incubated with an anti-mouse CD326 (EpCAM) AlexaFluor 647 antibody (1:100, clone G8.8, Biolegend) in 10% donkey serum in PBS for 3 days. Finally, the tissue was washed with PBS-T for 1 day.

### Quantification of crypts in whole-mounts

Imaging was done on a TCS SP5 confocal microscope (Leica). Images were analysed using Fiji. For SI, a minimum of 2,500 crypts per animal was scored. For colon, at least 900 crypts per mouse were scored. For the low-power analysis of clonal events, tdTom^+^ clones were scored across the entire length of the SI whole-mounts using a stereomicroscope (Nikon).

### Immunohistochemistry

For immunohistochemistry SI and colon were opened and fixed for 24 h in 10% neutral buffered formaldehyde in PBS. The tissue was paraffin embedded and sectioned by the CRUK CI Histopathology core. Haematoxylin and eosin staining was performed using an automated ST5020 Multistainer (Leica Biosystems). Alcian Blue and Periodic Acid/Schiff staining was carried out by the CI Histopathology Core. Briefly, slides were incubated in Alcian Blue for 10 min, and washed in water. They were then incubated in 0.5% periodic acid for 5 min, and washed 3 times. Slides were incubated in Schiff’s reagent for 15 min, washed 3 times, and counterstained with Mayer’s Haematoxylin.

BrdU and lysozyme immunohistochemistry was carried out using a Bond Max autostainer (Leica), with a proteinase K antigen retrieval. Slides were blocked with 3% hydrogen peroxide, followed by incubation in Avidin/Biotin Blocking Kit (Vector Laboratories). BrdU was detected using a sheep anti-BrdU antibody (1:500, Abcam ab1893). Rabbit anti-lysozyme antibody (1:500, Dako A0099) was used for lysozyme staining. Secondary antibodies in the two cases were biotinylated donkey anti-sheep (1:250, Jackson ImmunoResearch 713-066-147) and biotinylated donkey anti-rabbit (1:250, Jackson ImmunoResearch 711-065-152), respectively. Slides were incubated with Streptavidin coupled with horseradish peroxidase (HRP), and colour developed using diaminobenzidine (DAB) and DAB Enhancer (Leica).

Synaptophysin and Chromogranin A detection was carried out by manual IHC. Antigen retrieval was performed with 10 mM citrate buffer (pH 6.0) in a pressurised heating chamber. Tissue sections were incubated with rabbit anti-Chromogranin A antibody (1:500, Abcam ab15160), or rabbit anti-Synaptophysin antibody (1:300, Millipore AB9272), overnight at 4°C. Slides were incubated with biotinylated donkey anti-rabbit secondary antibody (1:500, Jackson ImmunoResearch 711-065-152). Streptavidin-HRP conjugate (Vector Laboratories) was added onto the slides and incubated for 30 min. DAB Chromogen substrate (Dako) was added for dye development. Counterstaining and dehydration was performed on the ST5020 Multistainer (Leica) followed by coverslipping.

### Immunofluorescence

For immunofluorescence tissue was excised and fixed for 48 h in 4% PFA in PBS at 4°C, after which it was transferred to 20% sucrose solution. After cryosectioning antigen retrieval where needed was accomplished by incubating the slides in 1% SDS for 5 min. Blocking was performed with 5% donkey serum. Following a wash primary antibodies were added and incubated overnight at 4°C. The following primary antibodies were used: rabbit FITC-anti-Lyz (1:400, Dako, F0372), rabbit anti-Muc2 (1:50, Santa Cruz, sc-15334), anti rabbit anti-ChgA (1:100, Abcam, ab15160). Secondary detection was with AlexaFluor 488 donkey anti-rabbit secondary antibody (1:500, Thermo Fisher). Alkaline phosphatase activity was detected using Blue AP kit (Vector Laboratories). Sections were covered with Prolong Gold with DAPI (Life Technologies). Fluorescent imaging was carried out on a TCS SP5 confocal microscope (Leica).

### Colon ulceration scoring

H&E-stained sections of colons were scanned on Aperio slide scanner (Leica Biosystems), and analysed using eSlide Manager (Leica Biosystems). Ulceration was defined as a region of a complete loss of crypt architecture and high cellularity.

### Tissue processing, smFISH and imaging

Harvested SI and colon tissues were flushed with cold 4% formaldehyde (FA) in PBS and incubated first in 4% FA/PBS for 3 hours, then in 30% sucrose in 4% FA/PBS overnight at 4°C with constant agitation. Fixed tissues were embedded in OCT. Quantification of co-expression was achieved by smFISH. Probe library design, hybridization procedures, and imaging settings were carried out according to published methods (Itzkovitz et al., 2011; Lyubimova et al., 2013). A Nikon-Ti-E inverted fluorescence microscope equipped with a Photometrics Pixis 1024 CCD camera was used to image a 10 μm cryo-section. A stack of 30 frames with 0.3 μm intervals was acquired to allow 3D cell imaging. FITC-conjugated antibody for E-cadherin was added to the hybridization mix and used to visualize cell borders. Detection of cells that were positive for Lgr5 transcripts, Atoh1 transcripts or both was performed manually with Fiji.

### Analysis of gut sections

Stained longitudinal sections of the SI and colon were visualised and positive cells scored manually. BrdU^+^ and negative nuclei were scored in complete half-crypt sections. Lysozyme^+^ cells were counted per whole crypt section. Alcian Blue/PAS^+^ cells were counted in complete half-villus sections, between the crypt neck and the tip of the villus. Cells in which the stain was clearly associated with a corresponding nucleus were marked as positive. Chromogranin A^+^ and synaptophysin^+^ cells were scored per complete half-crypt-villus section. Positive and negative crypts were scored, and results expressed as a frequency of positive cells.

### RNA isolation

For gene expression analysis by qPCR, cells were lysed and RNA isolated using RNeasy Mini Plus kit (Qiagen). For sequencing, total RNA was isolated from flow-sorted cells using RNeasy Micro Plus kit (Qiagen).

### Gene expression analysis

RNA was converted into cDNA (iScript cDNA synthesis kit, BioRad), and gene expression was analysed using TaqMan gene expression probes (Thermo Fisher). The following probes were used: Atoh1 (Mm00476035_s1), Muc2 (Hs00894053_g1), Tff3 (Hs00902278_m1), Spdef (Hs01026050_m1), Dll4 (Hs00184092_m1), Rassf4 (Hs00604698_m1), Gapdh (Hs02758991_g1).

### RNA sequencing

RNA quality was assessed on a 2100 Bioanalyser instrument (Agilent), according to the manufacturer’s instructions. The libraries were prepared using TruSeq Stranded mRNA Library Prep Kit (Illumina) and sequenced as 50 bp single-end reads on the Illumina HiSeq 4000 system.

### Western blotting

Protein extracts for SDS-PAGE were prepared by lysing the cells with RIPA buffer containing protease and phosphatase inhibitor cocktail (Thermo Fisher). Mouse anti-Atoh1 antibody (1:100, Developmental Studies Hybridoma Bank) and a rabbit anti-β-actin antibody (1:5,000, ab626, Abcam) were used. Fluorescent secondary antibodies were used (LiCor, goat anti-mouse 800LT (1:5,000), goat anti-rabbit 680LT (1:20,000)). For some experiments, protein extracts were incubated with λ phosphatase (New England Biolabs) prior to western blotting, according to the manufacturer’s instructions.

### *In vitro* kinase assay

The assay was performed as previously described(Azzarelli et al., 2017), with minor modifications. HA-tagged WT and mutant Atoh1 were *in vitro* translated (TNT® Quick Coupled Transcription/Translation Systems, Promega) in the presence of LiCl (800 mM) to reduce potential phosphorylation in reticulocyte lysate. Samples were incubated with human recombinant CDK/Cyclins (0.25 μM final concentration) in the presence of 10 μM ATP for 1 h at 30°C. Proteins were separated on Phos-tag™ gels (Alpha Laboratories, 7.5% acrylamide, 50 μM phos-tag PAGE, Wako) and immunoblotted with rat anti-HA-Peroxidase (1:5000, Roche).

### Cell proliferation and cell cycle analysis

Cell proliferation was assessed by an automated live-cell imaging system (IncuCyte ZOOM, Essen Bioscience). For cell cycle analysis, the cells were trypsinised, washed, fixed with ethanol, and stained with propidium iodide prior to flow cytometry.

### RNA sequencing analysis

The reads were aligned to the mouse reference genome [GRCm38] using TopHat2 aligner (Kim et al., 2013). Differentially expressed gene lists were generated using DESeq2 package from Bioconductor (Love et al., 2014).

### Secretory signature gene list

The list of differentially expressed genes (p<0.01) was generated by comparing the transcripts from tdTom^+^ and tdTom^-^ cells of *Atoh1*^*(WT)CreERT2*^ *Rosa26*^*tdTom*/+^ mice following tamoxifen. Upregulated genes in tdTom^+^ cells were selected to define a secretory signature in the small intestine and colon (Table S1). The top 500 upregulated, differentially expressed genes were used to perform the Gene Set Enrichment Analysis (GSEA).

### GSEA

This analysis was performed using the GSEA software from the Broad Institute (software.broadinstitute.org/gsea/index.jsp) (Subramanian et al., 2005). The list comprised all differentially expressed and non-differentially expressed genes from the *9S/T-A* v *WT* comparison in SI and colon, respectively. This gene list was probed with the previously generated secretory signatures (top 500 upregulated genes), and the published Atoh1^+^ gene signatures for ileum, colon (Lo et al., 2017), and intestinal stem cells (Muñoz et al., 2012).

### Data analysis

Statistical tests were not used to predetermine sample size. Randomisation was not performed to allocate samples/animals to experimental groups. Blinding was performed for quantifications in Figures 4A and 4B, as well as Figures S2A-S2E. Data analysis was performed using GraphPad Prism software or R package.

---

**Figure S1.**
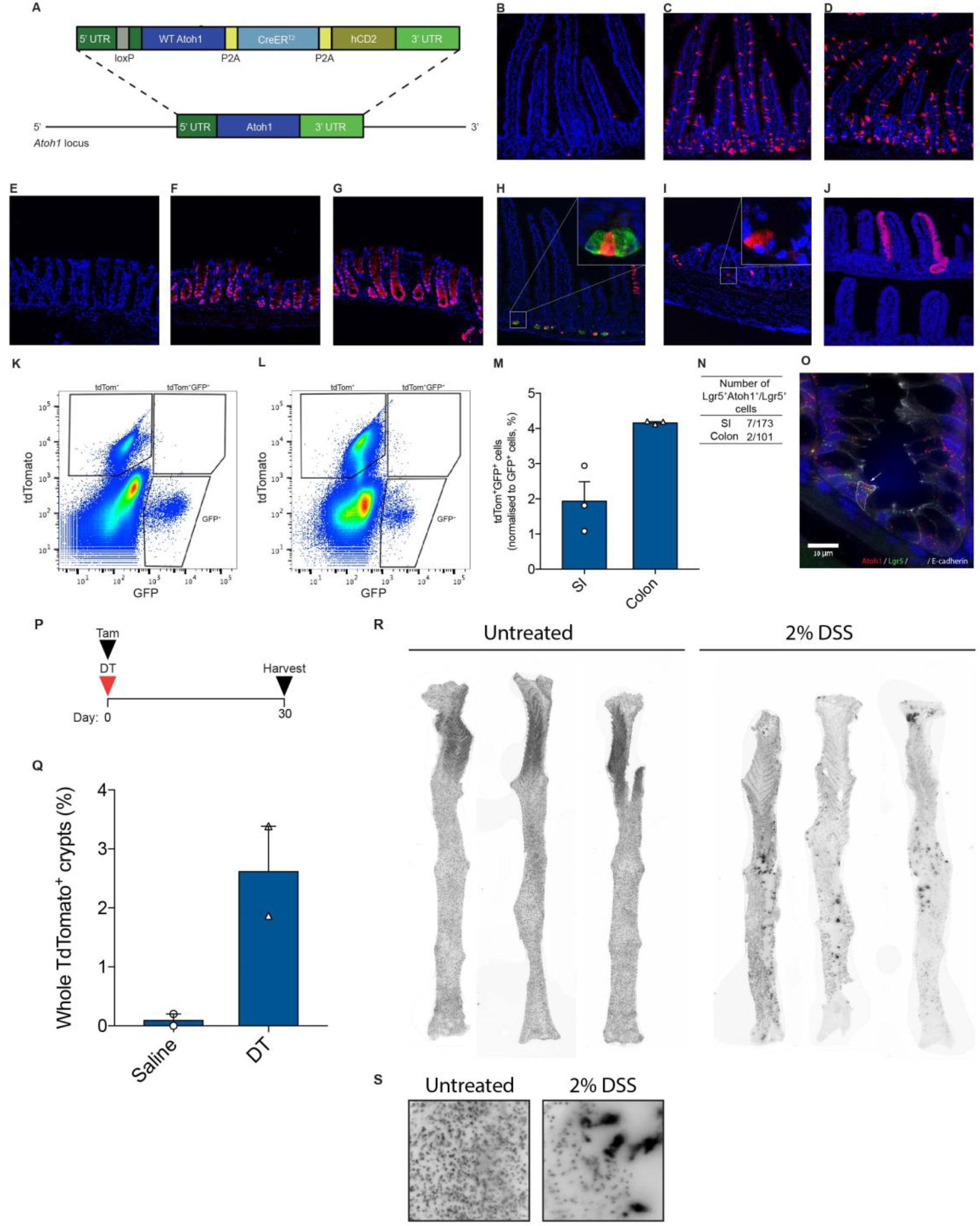
Generation of *Atoh1*^*(WT)CreERT2*^ mouse model and validation of lineage tracing in *R26R*^*tdTom*^ reporter line in homeostasis and injury, Related to Figure 1. (A) Schematic of the lineage tracing construct inserted into *Atoh1* genomic locus. (B, E) SI (B) and colon (E) of the uninduced animals. (C, F) Reporter positive cells in the SI (C) and colon (F), 24 h post-tamoxifen. (D, G) tdTom^+^ cells in the SI (D) and colon (G), 4 days post-tamoxifen. (H) Long-lived Lyz^+^ Paneth cells observed 30 days post-induction. (I) Long-lived reporter positive cells in the upper colon detected 30 days post-tamoxifen. (J) Labelled clonal event in the SI, 4 months post-tamoxifen. (K, L) Representative FACS graphs of tdTom^+^GFP^+^ cells in SI (K), and colon (L), 24 hours post-tamoxifen. (M) Quantification of tdTom^+^GFP^+^ cells in SI and colon. (N, O). Quantification (N) and a representative image (O) of Atoh1/Lgr5 transcript-positive cells in *WT* SI and colon (n=3). (P) Schematic of the induction/injury protocol. (Q) Quantification of tdTom^+^ clones in the SI without or with DT treatment (n=2 for each group). (R, S) Images of scanned colons showing tdTom^+^ crypts and multicrypt patches (MCPs) used for scoring in Figure 2T. DT, diphtheria toxin.

**Figure S2.**
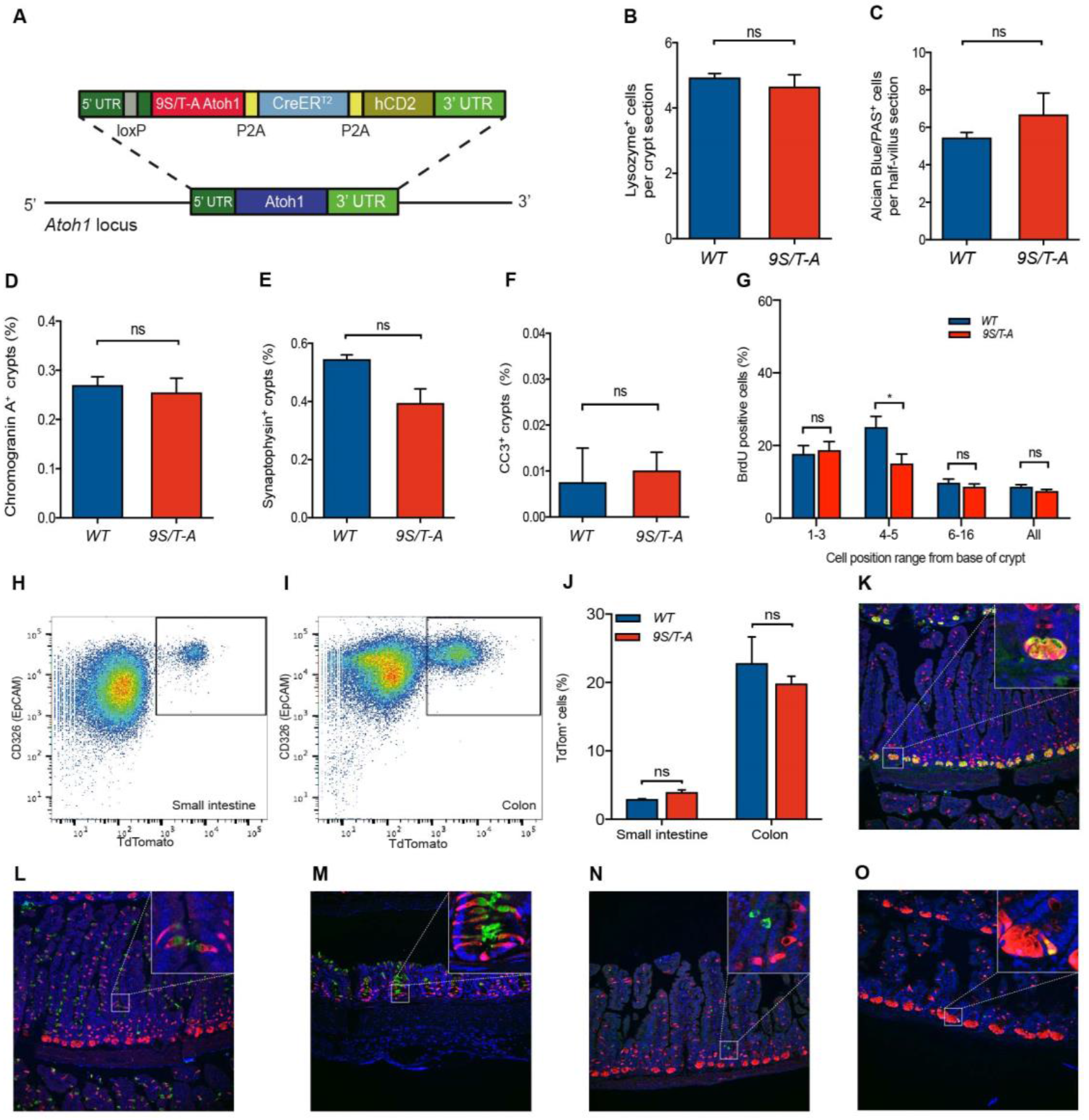
*Atoh1*^*(9S/T-A)CreERT2*^ mice do not exhibit abnormal intestinal secretory cell numbers and are phenotypically normal, Related to Figure 2. (A) Schematic representation of the *Atoh1*^*(9S/T-A)CreERT2*^ inserted into the *Atoh1* locus. (B-F) Quantification of differentiated small intestinal secretory lineage cells and apoptotic cells shows no difference between the genotypes. Paneth cells (B), goblet cells (C), enteroendocrine cells (D, E), and apoptotic cells (F) were scored (n=4 mice per group). (G) BrdU labelling index for a range of cell positions in colonic crypts (n=100 crypts, 4 mice per genotype; mean±s.e.m; * P=0.0101). (H, I) Representative flow cytometry plots of reporter positive cells in the SI (H) and colon (I). (J) Quantification of tdTom^+^ cells in the SI and colon of the two mouse lines (n=3 per group). (K-O) tdTom-labelling pattern in *9S/T-A* is identical to that of *WT*. Lyz^+^ Paneth cells (K), Muc2^+^ goblet cells in the SI (L), and colon (M) are positive for the reporter 24 h post-tamoxifen. ChgA^+^ enteroendocrine cells do not co-label with tdTom after 24 h (N), but acquire the reporter by day 4 post-induction (O).

**Figure S3.**
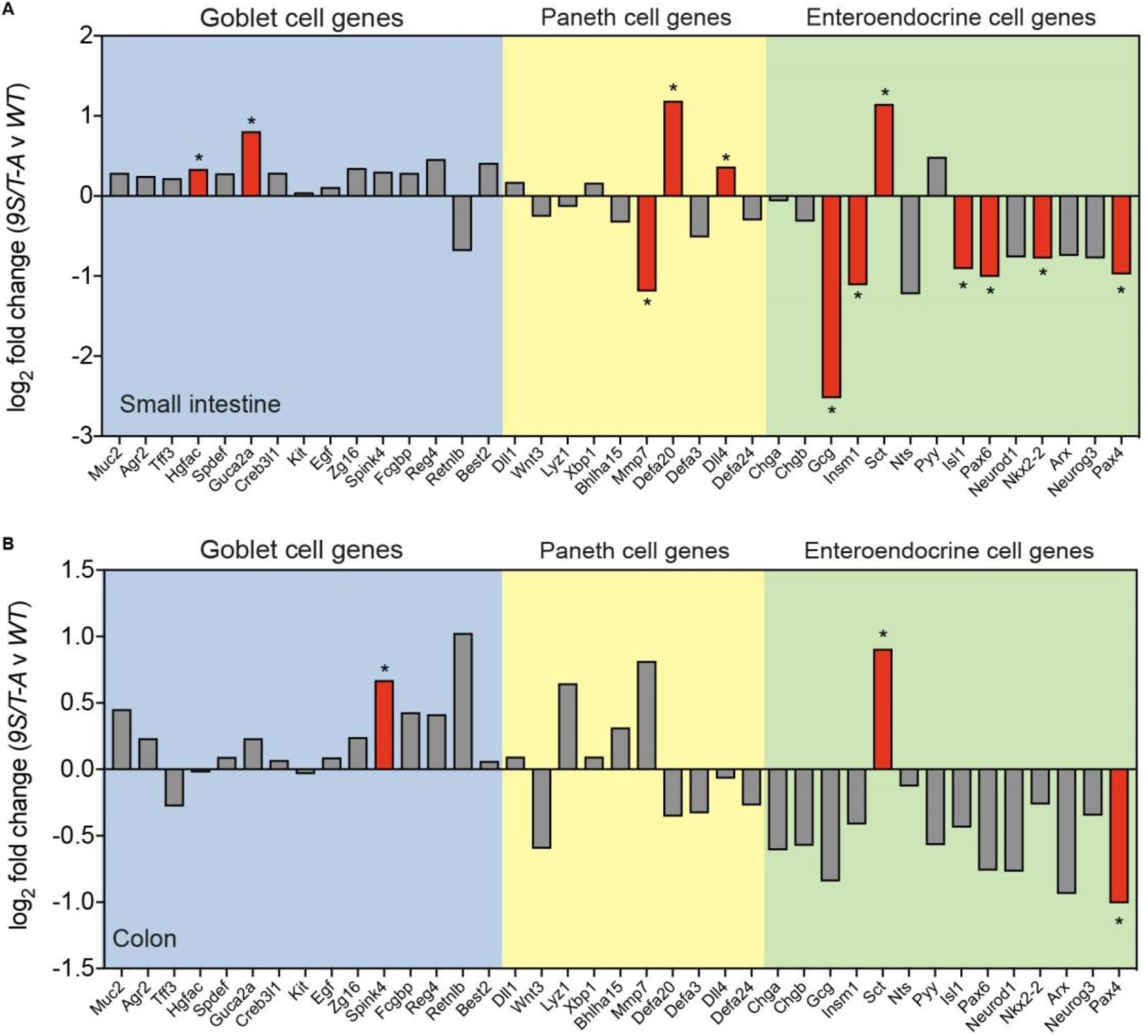
Secretory gene signature is upregulated in tdTom^+^ cells expressing 9S/T-A Atoh1, Related to Figure 3. (A, B) Expression of secretory genes selected based on known lineage-restricted pattern (Lo et al., 2017) in SI (A) and colon (B) of 9S/T-A Atoh1-expressing cells. Genes that are significantly differentially expressed (FDR < 0.1) are shown in red, labelled with an asterisk. The results were generated from n=6 for both *WT* and *9S/T-A* groups.

**Figure S4.**
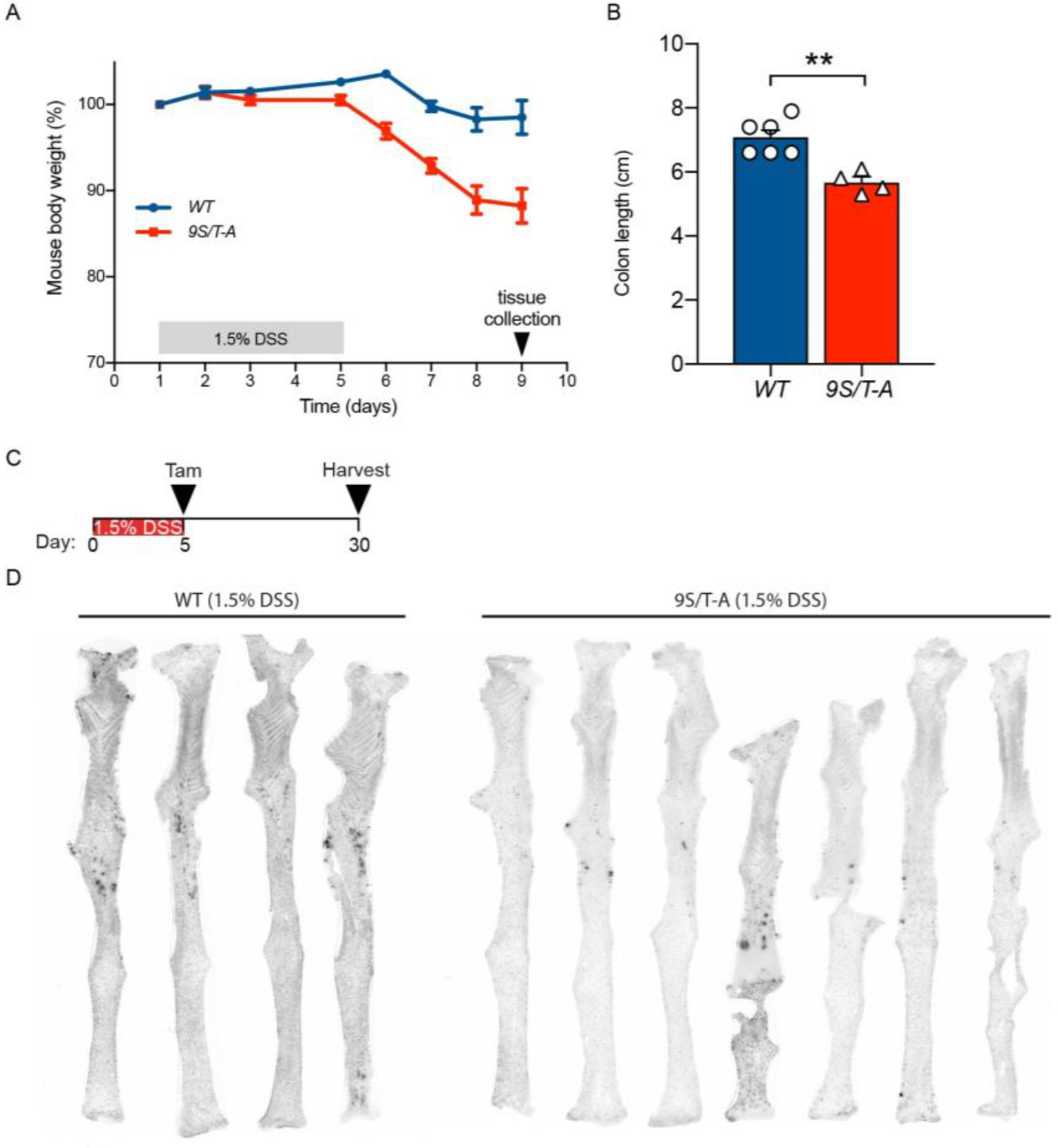
Characterisation of chemical colitis in *Atoh1*^*(9S/T-A)CreERT2*^ mice, Related to Figure 4. (A) Mouse body weight difference between *WT* and *9S/T-A* mice on day 9 (n=6 (*WT*), n=4 (*9S/T-A);* related to Figure 4B and 4C). (B) Colon length following DSS treatment (n=6 (*WT*), n=4 (*9S/T-A*), mean±s.e.m, ** P=0.0048). (C) Lineage tracing protocol following DSS colitis. (D) Images of scanned colons used for quantification in Figures 4G and 4H.

## SUPPLEMENTAL INFORMATION

### Computational analysis

The process by which crypt stem cells replace each occurs in a random though predictable manner. This behaviour can be modelled via a stochastic birth-death process (Lopez-Garcia et al., 2010; Snippert et al., 2010). The model was derived to model experiments where a single stem cell is labelled in a handful of crypts. As the number of initially labelled crypts was not of interest and to bypass any variability coming from the initial induction, make the different time points comparable, the equations were rescaled to account for only the surviving clones. Here we know the parameters of the stem cell dynamics (Kozar et al., 2013; Vermeulen et al., 2013), and would like to know the starting number of labelled stem cells per crypt and the number of labelled crypts.

For this analysis we use the equations described previously (Lopez-Garcia et al., 2010; Snippert et al., 2010), reproduced below. The probability of a crypt having clone of size n (for 0<n<N) at time t is:

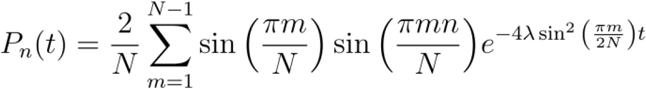

Here n is the number of labelled stem cells, N is the total number of stem cells, λ is the rate of stem cell replacement. And for the probability of all stem cells labelled we have:

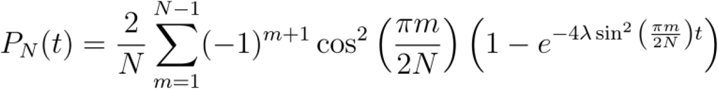

These equations assume the initial conditions of one labelled stem cell at t=0. The starting labelled stem cells were chosen randomly at the beginning of each simulation.

The values we observe for the clonal frequencies are substantially lower than what the model would predict, suggesting that not all crypts have labelled stem cells. In order to find out the fraction of labelled crypts *ν* we use a mixture model:

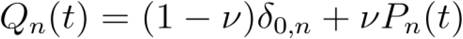

Where *Q*_*n*_(*t*) is the probability that a randomly selected crypt has a clone of size n labelled stem cells at time t. We use the values of N, λ and *τ* from Kozar et al and Vermeulen et al and estimate *ν*.

### Model fitting

For every mouse, at day 30 we count the number of clones (*k*_*i*_) and the number of crypts *C*_*i*_. We use a hierarchical model to capture the mouse to mouse variability. The statistical model is a follows

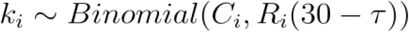

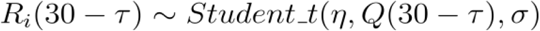

Here *R*_*i*_ is truncated to [0, 1]. For the SI no distinction is made in clone size, so Q is the sum of all *Q*_*n*_ and for the colon we use only the full clones for fitting *Q* = *Q*_*N*_.

The priors on the population parameters are:

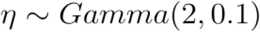

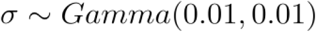

The prior on the mixing coefficient is

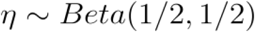

The posterior was derived via MCMC using Rstan (Carpenter et al., 2017). For the proximal and distal SI we used *τ* = 5 as the clones were measured in ribbons coming out of the crypt, which take a few days to emerge from the crypt base. Whereas for the colon we used *τ* = 1. The parameters used were λ = 0.1, *N* = 5 for proximal SI, λ = 0.2. *N* = 6 for distal and λ = 0.3, *N* = 7 for colon.

